# Bax Forms a Membrane Surface Protein-Lipid Complex as it Initiates Apoptosis

**DOI:** 10.1101/2022.11.01.514672

**Authors:** Luke A. Clifton, Hanna P. Wacklin-Knecht, Jörgen Ådén, Ameeq Ul Mushtaq, Tobias Sparrman, Gerhard Gröbner

## Abstract

Cellular clearance by apoptosis is essential in life. In its intrinsic (mitochondrial) pathway apoptotic members of the Bcl-2 (B cell CLL/lymphoma-2) protein family, such as Bax (Bcl-2-associated X) protein, perforate the mitochondrial outer membrane (MOM), which causes release of apoptotic factors and final cell death. How those apoptotic proteins mechanistically exert their action at the membrane level still however remains elusive. Upon internal stress signals Bax is massively recruited to the MOM, where it oligomerizes and partially penetrates into the membrane. Using neutron reflectometry (NR) and attenuated total reflection Fourier-Transform infrared spectroscopy (ATR-FTIR) spectroscopy we unraveled key molecular steps of this membrane-affiliation process of Bax on a spatial and temporal scale. By titrating intact human Bax to MOM-like bilayers containing cardiolipin, essential for protein recruitment, we could identify different functional phases. Initially, there is a fast adsorption event to the membrane surface with high affinity. Thereafter, a kinetically slower (minutes to hours) event occurs with Bax penetration, thereby triggering a major reorganizing of the mitochondrial bilayer. Finally, a membrane-Bax complex is generated, with a minor Bax population remaining membrane-inserted, while the main population is relocated to the membrane surface upon lipid redistribution into a complex with Bax; a process enabling membrane perforation. We propose a comprehensive molecular model of mitochondrial membrane penetration by formation of complex Bax/lipid clusters; a concept which provides a new foundation to understand the cell-killing activity of Bax and its apoptotic relatives in human cells.

**Significance Statement:** The apoptotic Bax protein is a key player in the mitochondrial apoptotic pathway. Here, neutron reflectometry (NR) unravels the mechanism by which Bax is targeting and perforating mitochondria to release apoptotic factors for final cell death. We found that this cardiolipin driven process of the outer mitochondrial membrane system has two main phases. Upon a fast (10-20 min) phase of membrane association Bax initiates the formation of pores by removing lipids and depositing them as Bax/lipid complexes on top of the bilayer on a time scale of several hours similar to *in vivo* apoptotic cell death. Our results provide a mechanistic rationale for cell-killing processes driven by apoptotic Bcl-2 proteins; and their molecular inhibition in many cancers.

## Introduction

Programmed mammalian cell death, also denoted apoptosis, is essential for embryonic development, tissue homeostasis and removal of harmful cells (1, 2). Failure in its regulation can cause various pathological disorders including cancer (3, 4). Upon activation of the intrinsic apoptotic pathway by intracellular stress, mitochondria become intimately involved in its further progression towards cellular death (5). During this process the mitochondrial shell – its mitochondrial outer membrane (MOM) - undergoes permeabilization, thereby enabling release of apoptotic factors such as cytochrome which triggers an irreversible signaling cascade (6). To protect healthy cells from undesired clearance, this pathway is tightly regulated by the Bcl-2 (B-cell lymphoma 2) protein family (5, 7-9). Pro- and anti-apoptotic members of this family meet at the MOM and they arbitrate the cell’s fate there: intact membrane and survival *versus* permeabilization and death (1, 2). Failure in this regulation process can cause various pathological disorders ranging from embryonal defects to cancer (3, 4).

The most prominent multidomain members of the Bcl-2 family are the apoptotic Bax and the anti-apoptotic Bcl-2 protein itself. Since those proteins have to be membrane-active to interact at the MOM and control its integrity, they possess the typical features of “tail anchored membrane proteins” with an amphitropic main globular fold and a single transmembrane (TM) domain to enable association to and penetration into the MOM target membrane (5, 6). While Bcl-2 is insoluble with its TM domain stretched out to anchor and immerse the protein into its membrane environment (7), Bax is soluble by hiding its TM domain inside its hydrophobic groove region (8). Normally, Bax is mainly found in the cytosol, but upon activation by apoptotic stimuli it can expose its TM domain in order to be recruited to the MOM surface and, partially penetrate into it and finally perforate it to subsequently release apoptotic factors (9-11). In healthy cells, this pore formation ability by Bax is prevented by its tight binding to the pro-survival Bcl-2 protein which is located at the MOM. However, upon apoptotic stimuli Bax is massively recruited to the MOM where its unique lipid cardiolipin (CL) playing a driving role (12, 13). Subsequently it undergoes conformational changes and finally forms dimers and larger oligomers causing membrane pore formation and lethal release of apoptotic factors (14, 15). However, high-resolution structural information about membrane-bound Bax and its pores is lacking, despite recent progress on pore characterization by computational approaches (15), and by AFM and fluorescence based advanced studies (14, 16, 17). Therefore, the precise mechanism driving Bax membrane association and assembly into membrane pores remains quite unclear at the molecular level for all components involving the protein itself and to the MOM lipid matrix.

To obtain a molecular understanding of the ability of the Bax protein to recognize mitochondrial membranes upon activation and to permeabilize them, we studied the spatial and temporal organization of intact human Bax protein and membrane lipids during this process using MOM-like supported bilayers with varying amounts of cardiolipin, a mitochondria-specific lipid essential for membrane association of Bax, its activation there and the subsequent pore formation during apoptosis (12, 18). To follow the locations, including time-dependent displacements, of the individual components in the membrane/Bax complexes we applied two complementary approaches: neutron reflectometry (NR) on supported lipid bilayers and attenuated total reflection Fourier-Transform infrared spectroscopy (ATR-FTIR) spectroscopy. During recent years, NR with isotopic contrast variation has become a powerful technique for providing a general overview of the structure across membrane systems as it allows the relative distribution of individual components in these complex structures to be resolved (19-21). Upon titration of Bax to various supported bilayer systems, NR enabled us to monitor the reorganization of Bax and lipid components as a function of time and membrane location by evaluating their relative distributions across the bilayer and visualizing them as scattering length density (7) and corresponding component volume fraction profiles (22). In addition, NR can also provide adsorption kinetics of molecules onto surfaces of different types. For this purpose, time-resolved NR data were acquired by us to monitor Bax adsorption onto lipid bilayers and the subsequent ongoing changes of the entire Bax/membrane assembly. This type of spatial and temporal information on a global scale for molecular systems is otherwise nearly impossible to access by atomic-level approaches such as NMR spectroscopy or x-ray crystallography. As complementary method, ATR-FTIR provides valuable information to study changes in chemical composition of supported bilayers. Since chemical groups have specific absorption bands, different chemical groups in a sample can be monitored simultaneously, and besides sample composition, it provides information about time-dependent changes in composition, secondary structural changes of proteins, lipid phase transitions and more (23). Here, we used protein-specific IR bands (amide-I band) and the lipid acyl chain (CH) stretches to monitor the kinetic and structural changes of Bax upon titration onto lipid bilayers. The outcome of the time-dependent NR and ATR-FTIR experiments on Bax titration series onto MOM-mimicking bilayers with varying CL content under varying isotope contrast conditions revealed three major distinct events. Upon a rapid initial adsorption onto the target membrane, activated Bax penetrates further into the membrane on a slower timescale causing finally its perforation by depositing Bax/lipid assembly onto the bilayer. The entire event occurs on a time scale of 2-5 h which is of the same order of magnitude as *in vivo* apoptosis (24). The detailed molecular mechanistic picture obtained by our time-resolved NR approach provides a solid foundation to understand the membrane-destroying function of apoptotic multi-domain members of the Bcl-2 family in general and opens up the way for overcoming their molecular inhibition in many tumors in the search for novel cancer therapies.

## Results and Discussion

### Reorganization of the mitochondrial outer membrane in the presence of Bax

Neutron reflectometry (NR) enabled us to reveal the final stages of perturbations occurring in MOM-like bilayers in the presence of intact human Bax protein as seen in Fig. 1. Vesicle fusion was used to deposit a series of supported lipid bilayers composed of a mixture of the zwitterionic phospholipid POPC and the anionic phospholipid TOCL, with this mixture acting as representation of the lipids found in the mitochondrial outer membrane (25). Three individual mixtures of POPC and TOCL were studied with increasing CL content (5 mol%, 10 mol% and 15 mol%), which represents the heterogenous CL distribution across the MOM (25). In all cases structural analysis by NR revealed high surface membrane coverages approaching 100%. The interaction of Bax with the models of the mitochondrial outer membrane was examined applying NR for all three CL concentrations. Using 10 mol% CL containing bilayers as an example, Fig. 1 shows the corresponding NR data, model data fits and the scattering length density profiles before and upon the interaction of Bax under steady-state conditions (several hours) at 37°C. NR data reveals the structural changes and redistribution of lipids and protein components after the interaction of Bax with the model mitochondrial outer membrane.

**Figure 1.**
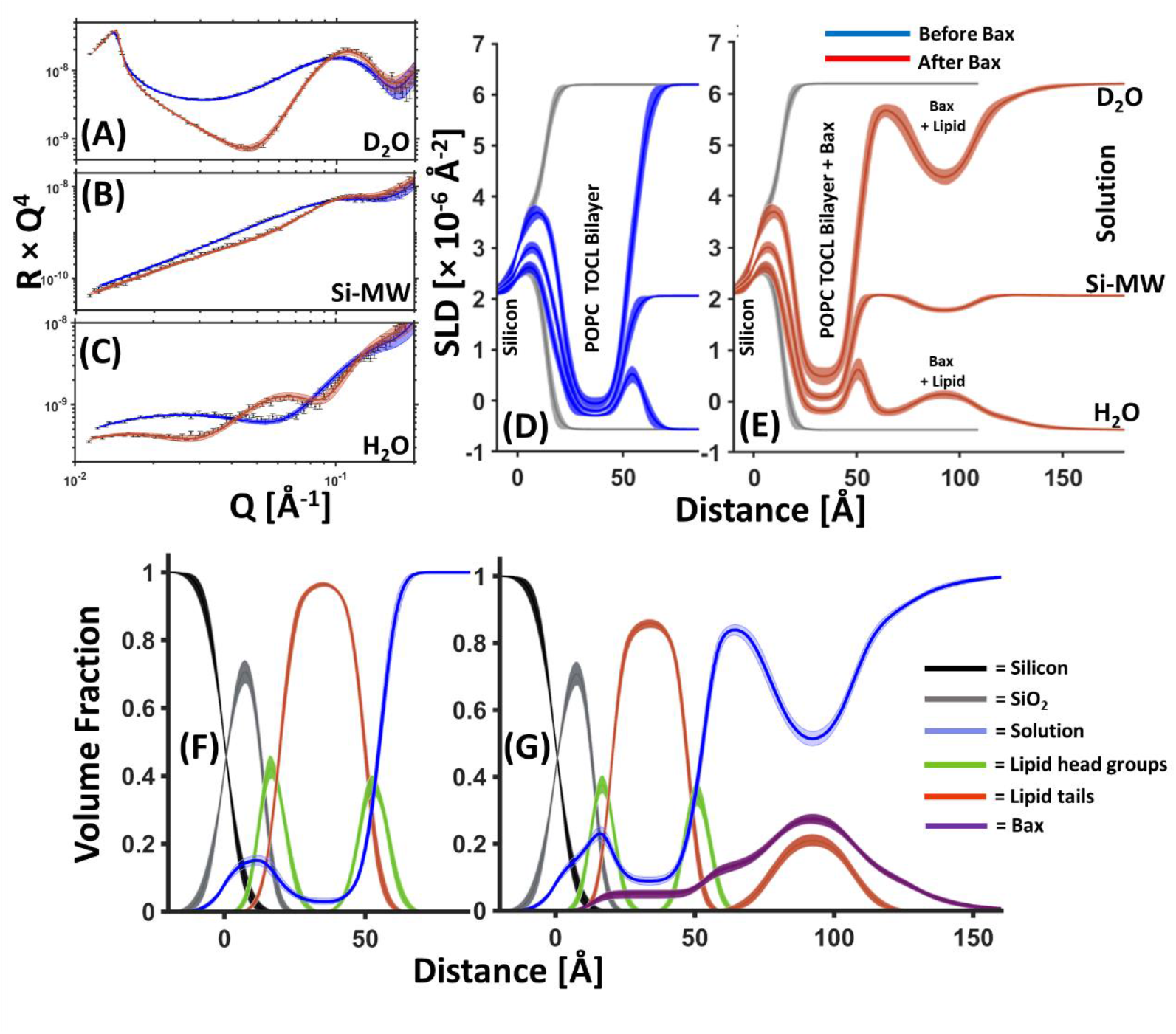
NR data showing Bax-induced redistribution of lipid from the membrane core to a surface protein-lipid complex during its apoptotic mechanism: NR data (error bars) and model data fits (lines; s. also supplement Table 1) from a 90 mol% POPC – 10 mol% TOCL SLB before (blue) and after (red) the interaction of natural abundance hydrogen (h-)Bax are shown in three differing solution isotopic contrast conditions being D_2_O (A), Si-MW (B) and H_2_O (C) buffer solutions. The scattering length density (SLD) profiles and the model fits are shown for the surface structure before (D) and after the h-Bax interaction (E). The corresponding component volume fraction profiles are shown before (F) and after (G) the h-Bax interaction as determined from the NR fits. Individual components are color-coded as indicated, with the Bax protein distribution in purple. Note that after the interaction of the protein there is a lower lipid content in the SLB and a new distribution of lipid on the membrane surface. Line widths in the NR data fits represent the 65% confidence interval of the range of acceptable fits determined from Monte-Carlo-Markov Chain (MCMC) error analysis and the line widths in the SLD and volume fraction profiles represent the ambiguity in the resolved interfacial structure determined from these.

Prior to the interaction of Bax the coverage of the MOM-like bilayers was 98±1 vol% lipid (see Table 1), after the interaction of the protein the lipid content of the SLB had been reduced to 86±2 vol% with an increase in the water content of the lipid tail region of the SLB from 2±1 vol% to 9±1 vol%, with 5±2 vol% protein also appearing in this region (Fig 1*F* and 1*G*). In addition, the SLB was also seen to become thinner with the SLB tail thickness reducing from ∼30 Å to ∼28 Å. Together with the observed increase in water and protein within the SLB these data are suggestive of the formation of pores within the membrane, an activity which is directly related to Bax’s role as an initiator of the biochemical stages of apoptosis (14, 16, 26).

**Table 1.** The resolved structural components before and after the interaction of Bax with SLB composed of POPC-10 mol% TOCL.*

|  | Average Lipid Area per Molecule [Å<sup>2</sup>] | Tails Thickness [Å] | Tails Composition [vol%] | Head Group Thickness [Å] | Outer head-group Composition [vol%] | Membrane Surface Bax/Lipid Complex Thickness [Å] | Bax Surface Layer Composition [vol%] |
| --- | --- | --- | --- | --- | --- | --- | --- |
| **POPC: CL (90:10 mol%)** | 64.5 Å<sup>2</sup><br>(63.8 Å<sup>2</sup>, 65.2 Å<sup>2</sup>) | 29.8 Å<br>(29.4 Å, 30.1 Å) | Lipid 98%<br>(97%, 99%)<br>Solution 2%<br>(1%, 3%) | 5.7 Å<br>(5.4 Å, 6.4 Å) | Head-group 83%<br>(81%, 89%)<br>Water 17%<br>(14%, 21%) | - | - |
| **POPC: CL (90:10 mol%) + h-Bax** | 69.7 Å<sup>2</sup><br>(68.7 Å<sup>2</sup>, 70.2 Å<sup>2</sup>) | 27.6 Å<br>(26.9 Å, 29.0 Å) | Lipid 86%<br>(85%, 88%)<br>Protein 5%<br>(3%, 6%)<br>Solution 9%<br>(8%, 10%) | 6.5 Å<br>(6.2 Å, 6.6 Å) | Lipid 65%<br>(63%, 68%)<br>Protein 5%<br>(3%, 6%)<br>Solution 30%<br>(27%, 34%) | **1**, 23.4 Å<br>(22.2 Å, 24.9 Å)<br>**2**, 29.7 Å<br>(27.1 Å, 31.2 Å)<br>**3**, 23.4 Å<br>(22.2 Å, 24.9 Å)<br>**Total : 76.4 Å**<br>**(75.2 Å, 77.7 Å)** | **Protein Lipid Distribution Composition:**<br>1, Protein 12% (10%, 14%)<br>Water 88 % (86%, 90%)<br>2, Protein 30 % (27%, 32%)<br>Lipid 17% (16%, 18%)<br>Solution 53% (51%, 55%)<br>3, Protein 12% (10%, 14%)<br>Water 88 % (86%, 90%) |

\* Values in brackets represent the 65% confidence intervals determined from MCMC resampling of the experimental data fits

The interaction with Bax produced a symmetrical protein-lipid complex on the membrane surface. The structure was composed of a complex layer of protein and lipid between two layers of protein. The amount of lipid within the membrane surface complex was optimally fitted to exactly match the amount of lipid removed from the SLB both through pore formation and membrane thinning. This result suggests that Bax is able to redistribute lipids from the membrane interior to the membrane surface in its pore forming activity.

Those changes in the interfacial membrane structures are representative for all samples with varying CL content. Measurements and corresponding analysis plots for SLBs containing 5 mol%, and 15 mol % CL are shown in supporting materials (*SI Appendix*, Fig. S1, S2). There was a noticeable trend in the NR data where the amount of lipid removed from the SLB to form pores was proportional to the initial CL content (see Table 2). Nevertheless, Bax induced membrane thinning was fairly consistent across all samples with a ∼2-3 Å decrease in bilayer thickness observed at equilibrium Bax binding (Table 1 and *SI Appendix*, Tables 1, 2 and 3).

**Table 2.** The relationship between SLB TOCL concentration and lipid loss during Bax driven membrane disruption.*

| **SLB TOCL Content** | **5 mol%** | **10 mol%** | **15 mol%** |
| --- | --- | --- | --- |
| **SLB Tails Coverage Before Bax / vol%** | 99%<br>(98%, 100%) | 98%<br>(97%, 99%) | 99%<br>(98%, 100%) |
| **SLB Tails Coverage After Bax / vol%** | 91%<br>(90%, 92%) | 86%<br>(85%, 88%) | 81%<br>(80%, 82%) |
| **Lipid Removed Through Pore Formation / vol%** | 8%<br>(6%, 10%) | 12%<br>(9%, 14%) | 18%<br>(16%, 20%) |

\*Tables of NR resolved structural parameters from the interaction of Bax with SLBs containing 5% and 15% are given in supporting materials Tables 1 and 2. Values in brackets represent the 65% confidence intervals determined from MCMC resampling of the experimental data fits.

### Schematic model of membrane perforation by Bax

As the mechanism of redistribution of the lipid from the membrane to the membrane surface by Bax is a novel finding, NR measurements on the interaction of Bax with a POPC-10 mol% TOCL SLB were repeated using deuterium labelled protein. The advantage of using deuterated (d-)Bax is that the d-protein will be contrast matched (i.e. invisible) in the D_2_O contrast condition (7) giving a data set which only describes the structure of the SLB and the lipid components. This data (*SI Appendix*, Fig. 3) provided independent confirmation that the interaction of Bax with the model membrane caused a redistribution of lipids from the SLB to the membrane surface, and again showed that the lipid component moved to the membrane surface matched the amount lost from the SLB during pore formation and bilayer thinning.

The thickness profile of the membrane-surface protein-lipid complex was suggestive of a structure which was three times the width of Bax in its solution folded state (27). We have proposed a model of the SLB surface structure (Fig. 2) where Bax forms a toroidal-like pore within the SLB, around which there is a protein lipid complex composed of three layers with one protein-only layer bound both between the SLB and the protein-lipid complex and another on top of the complex. Bax seems to perforate the MOM by sequestering membrane lipid components into Bax/lipid clusters deposited above the bilayer to make space for the pores. The formation of this membrane surface structure could be related to the ring structures found to enclose Bax-induced pores in previous atomic force microscopy studies (14). In that study activated Bax formed protruding Bax clusters (like coronas) around 4 nm above the SLB; a scenario well in agreement with our Bax-lipid complexes and their thickness distribution as seen in Fig. 2. The lipids excluded to create pores form proteolipid clusters above the SLB. In our model Bax forms pores at the membrane level with a low fraction of Bax residing inside the membrane level compared to that found on the membrane surface (s. Fig. 2*B* and *SI Appendix*, Tables 1, 2 and 3) which is partially in agreement with previous lipidic pore models (26, 28). The model clearly supports and fundamentally expands recent super-resolution microscopy and Cryo-EM studies showing in human cell formation of GFP-labelled Bax clusters with higher molecular organization near mitochondria (29). Even for Bak a close relative of Bax, disordered clusters of Bak dimers were found to generate lipidic pores (30). However, information about any lipid contribution to the Bax-mediated apoptotic steps could not be obtained by their approaches.

**Figure 2.**
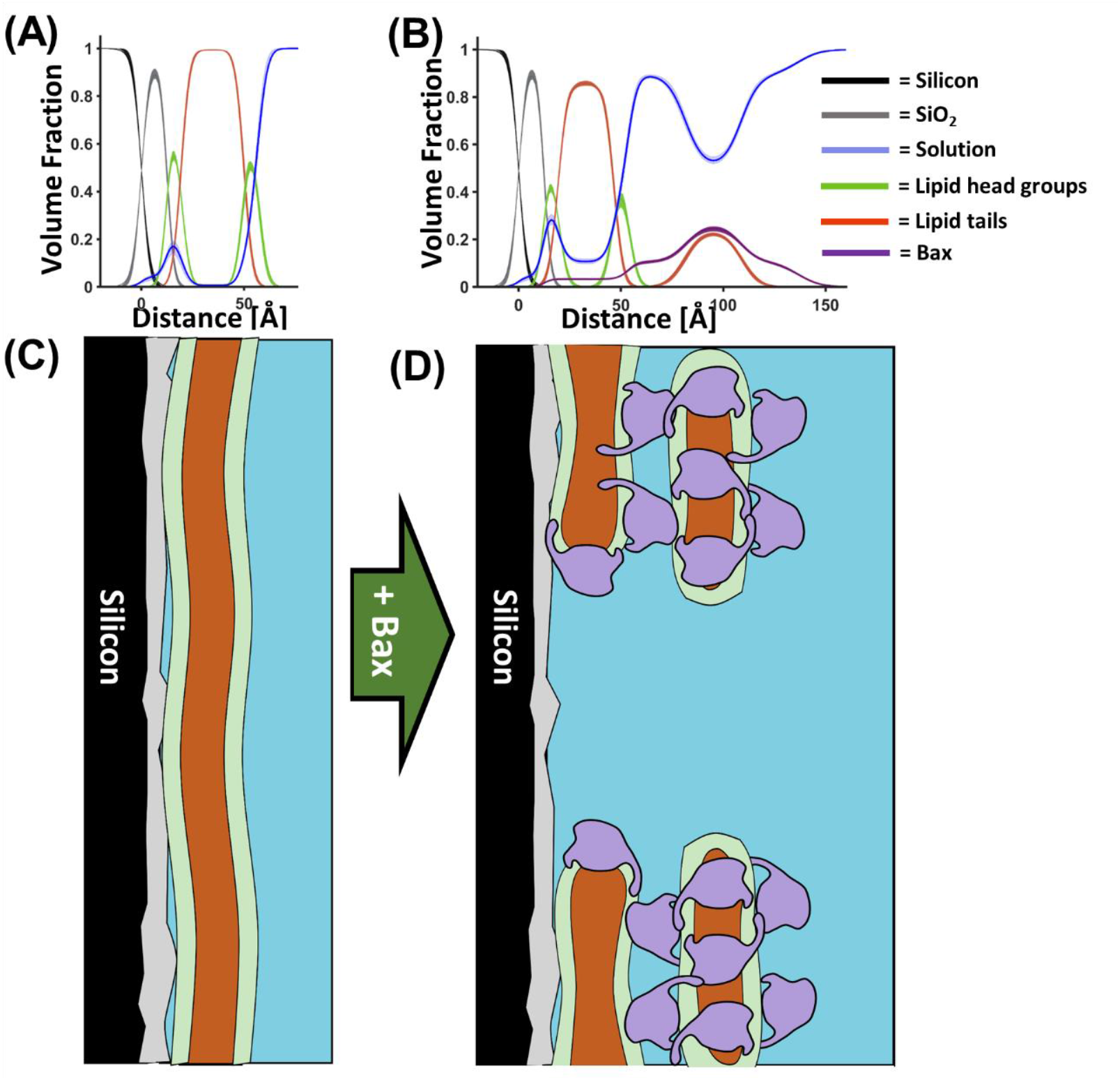
A model for Bax pore formation: Component volume fraction profiles from before (A) and after (B) the interaction of Bax with mitochondrial model membranes (in this case d-Bax to a 90 mol% POPC – 10 mol% TOCL SLB; see *SI Appendix*, Fig. 3). Below these are schematics of how this component distribution in the Z direction may relate to Bax mediated pore formation. This model comes from consideration of the data presented here and in previous SLB studies using AFM (14).

### Real-time kinetics of Bax induced permeabilization of mitochondrial outer membranes

In general, to create pores, Bax and even Bak have to undergo structural changes upon membrane contact to form dimers, which then oligomerize into protein clusters to perforate the membrane (16, 27, 29, 30). To in depth unravel the molecular principles and determinants by which Bax executes this complex process, in particular recognition and reorganization of the MOM, for enabling further apoptotic steps, we carried out a series of experiments to study kinetics and dynamics of Bax interactions and their dependence on the amount of cardiolipin in the target membranes. Therefore, a combination of NR and ATR-FTIR was used to give time dependent structural and mechanistic insight into the kinetics of the Bax/SLB interaction.

Monitoring the interaction of deuterated (d-)Bax with a fully hydrogenous (h-)membrane in a H_2_O buffer solution allowed collection of time-dependent structural data which was highly sensitive to changes in the protein distribution across the surface. This data, shown in Fig. 3*A*, was optimally fitted to show an increasing surface coverage of the full, three-layer, protein-lipid complex on the SLB surface against time (Fig. 3*B*). Independent measurements by ATR-FTIR probed the increase in the protein Amide I band of Bax against time (Fig. 3*C*), which gave an independent measurement of the accumulation of protein within the surface region of a SLB model of the mitochondrial outer membrane (7, 31, 32). A comparison between the changes in the volume fraction of the protein-lipid membrane surface complex and the increase in the Amide I band against time was favourable (Fig. 3*D*) suggesting two processes on different time scales. First, an initial rapid adsorption and accumulation of Bax on the membrane surface which decreases over time, similar to Langmuir isothermal adsorption behaviour; and second, a slower protein-lipid complex formation and perforation process. Fitting of the FTIR-data with a two-exponential kinetic model (blue line in Fig. 3*D*) provided a 10 +/-2 min time constant for the fast process and ca. 175 +/-15 min for the slower process. Analysis the NR data (red line) data for the initial fast process could not be obtained due to the restrictions in time resolution of the NR technique, but a time constant of 110 +/-30 min was obtained for the slower process; a value in quite good agreement with the FTIR results despite different sample environments and sample batches. Even studies using fluorescence labelled Bax overexpressed in intact cells observed heterogenous Bax cluster formation near mitochondria on a 100 min +/-60 min scale upon initial Bax recruitment to mitochondria (29).

**Figure 3.**
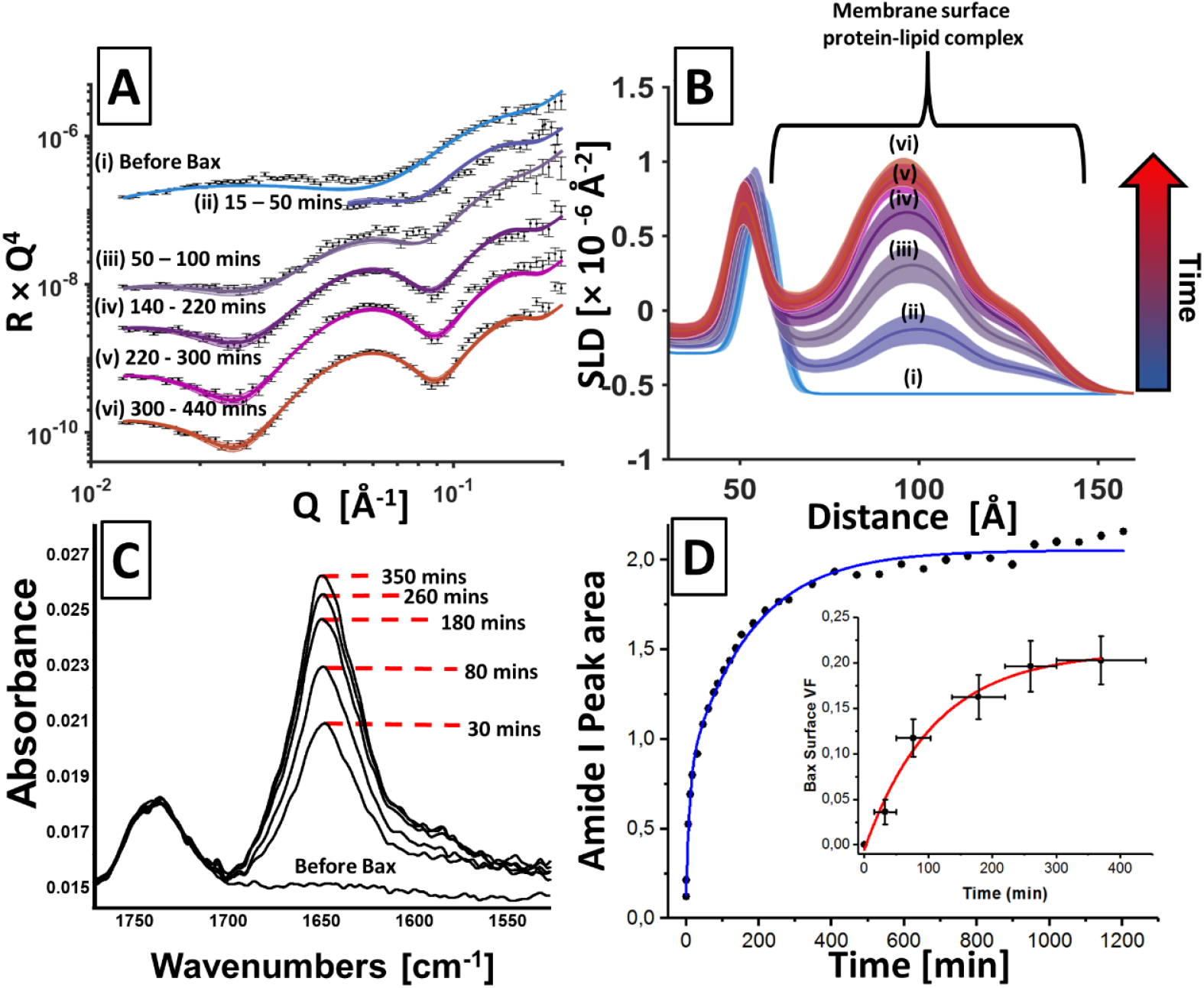
Time-dependent structural data based on the interaction of Bax with model mitochondrial outer membranes. NR data (error bars) and model data fits (lines) obtained in a H_2_O buffer from a POPC 10 mol % TOCL SLB before and at different times after introduction of deuterated (d-)Bax into the solid liquid cell containing the bilayer (A). The SLD profiles obtained from model fitting of this data showing the increasing coverage of Bax on the membrane surface with increasing time (B). Complementary ATR-FTIR measurements on the same experimental system show the increasing size of the Amide I band indicating the accumulation of Bax at the membrane containing solid-liquid interface (C). Finally, a comparison of the proteins surface volume fraction (crosses) and amide I peak areas (red) vs time obtained from NR and ATR-FTIR measurements respectively showing the strong agreement between in the two techniques on the kinetics of the Bax/membrane interaction (D).

Combined, these data suggest that the formation of the complex protein-lipid surface assemblies is commensurate with the Bax interaction with the SLB rather than a stepwise build-up of the three protein-containing layers on the membrane surface. Qualitatively, the two processes identified by us here directly by NR reflect earlier studies that suggest that in the first phase, Bax becomes activated upon initial membrane contact and partially inserts to generate dimers under structural rearrangements. Those dimers are then a prerequisite for forming supramolecular pore structures of Bax at the MOM on a slower time scale to initiate further apoptotic activities (14, 28, 33). This later process occurs during several hours as monitored in our NR experiments in real-time which agrees well with times reported for *in vivo* cell death to happen (24, 34): early apoptosis – a few hours; late apoptosis – 6 to 24 h.

### Role of Cardiolipin in driving Bax mediated membrane activities

The mitochondria-specific lipid cardiolipin (CL) plays a key role during apoptosis in facilitating recruitment of Bax to the MOM, the subsequent insertion of Bax into the membrane and its final perforation (10, 12, 18, 35). The distribution of CL across the MOM is quite heterogenous with an increase of CL (>20 mol%) near contact sites and a general increase during apoptosis (25, 36). Based on previous work on Bax-induced membrane leakage that occurs in a CL-dependent manner, we were interested in studying the time dependence of Bax/lipid assembly formation and outcome of membrane reorganization for MOM bilayers with varying CL content.

Therefore, time-resolved NR data was used to examine the relationship between the pore and membrane surface protein-lipid complex formation and the concentration of cardiolipin in the mitochondrial outer membrane model. Fig. 4*A* and 4*B* show time-resolved NR data obtained during the interaction of (h-)Bax with the 15 mol% TOCL containing mitochondrial outer membrane model obtained in a D_2_O buffer solution. NR data sets obtained in D_2_O are sensitive to both the lipid and h-protein distributions across the surface (37). This data was optimally fitted to models which were a scale ratio between the surface structure before the interaction of Bax and the structure at equilibrium Bax binding. This produced time-dependent structural changes where lipid removal from the SLB and membrane surface protein-lipid complex formation were proportional. This suggest either individual pores in the SLB grow larger over time or that the number of pores increases. The correlation between pore and surface complex formation gives further evidence that the membrane surface protein-lipid complex is a by-product of pore formation.

**Figure 4.**
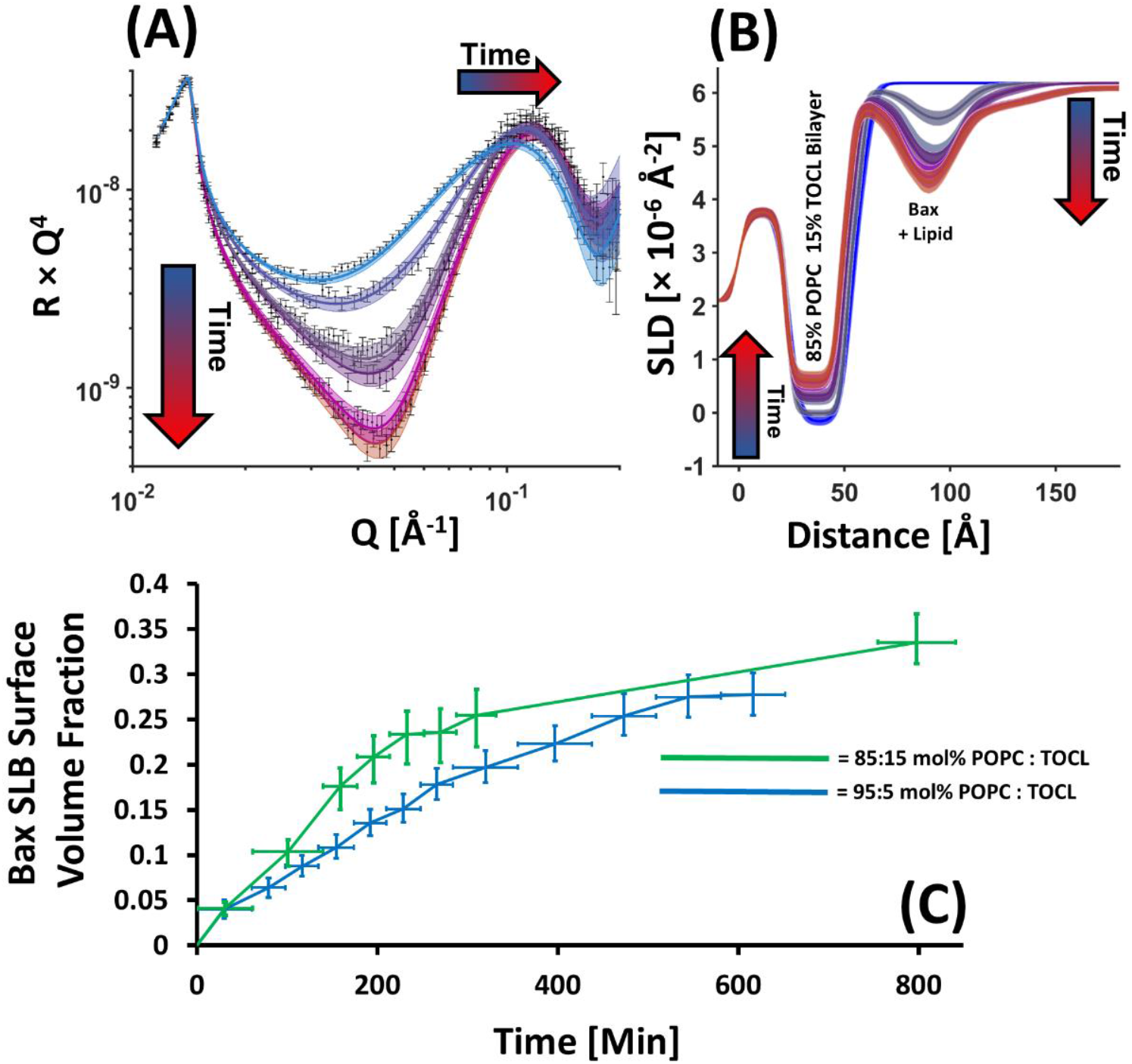
Time-resolved NR data revealed that pore formation and membrane surface protein-lipid complex formation increase proportionally with time and that the kinetics of the membrane reorganisation process is positively correlated to bilayer cardiolipin concentration. Time resolved NR data (error bars) and model data fits (lines) from an POPC-15 mol% TOCL SLB during the interaction of (h-)Bax in a D_2_O buffer solution (A) and the changes in the scattering length density of the surface assemblage against time (B, depicted from blue to red). Note how the increasing SLD of the bilayer, depicting lipid loss, is commensurate with decreasing SLD of the Bax/Lipid surface complex, which depicting increasing complex coverage. Lines widths on the fits/NSLD profiles depict the ambiguity in the resolved structure from MCMC error estimation. A comparison of the change in protein coverage on the membrane surface (from the highest density middle layer) from NR data from a series of mitochondrial outer membrane models with differing cardiolipin content (C). Each data point on this plot represents an individual NR data set, the error bars on the x-axis represent the range of time over which the data set was collected while the y-errors represent the ambiguity on protein volume fraction determined by MCMC resampling of the model-to-data fits. Time resolved NR data and fits for the interaction of Bax with the POPC-5 mol% TOCL bilayer are given in *SI Appendix*, Fig. 4.

The time dependence on the Bax interaction with the SLBs appeared to be correlated to the concentration of cardiolipin in the membrane. Fig. 4*C* shows a comparison of the change in membrane surface Bax volume fraction (from the highest density middle layer of the complex) from the interaction of the protein with the 15 mol% and the 5 mol% TOCL bilayers. This data suggests that the rate of interaction of Bax with the membrane is cardiolipin dependent. However, it was interesting to note that the final coverage of Bax on the membrane surface was similar in all measurements.

Despite a clear prerequisite for CL in the MOM for of Bax to be attracted to the membrane and perforate it (10, 12, 18, 35), we cannot distinguish any CL preference in the mechanism of pore formation that involves the removal of all lipid species (PC and CL) into proteolipid complexes (see Fig. 3, Fig. 4 and Table II). This indicates that, after partial penetration of Bax into CL containing membranes followed by pore formation, the formation of oligomeric Bax/lipid aggregates is not lipid specific but reflects the average lipid composition of the bilayer. From a biological perspective it could be important that CL gets removed into Bax/lipid clusters outside the mitochondrial membrane system to enable quick release of cytochrome c into the cytosol to initiate further irreversible apoptotic steps (12, 15), without a high probability to sequester again into CL/cytochrome c complexes. Further progress of the apoptotic pathway *in vivo* involves additional steps including cytochrome c release upon dissociation from CL at the inner mitochondrial membrane, followed by major redistribution of CL across the entire mitochondria including dramatic morphological changes of the mitochondria (29, 38, 39).

In summary, our NR experiments provide a molecular model and explanation of the overlapping events of pore formation and lipid complexation reflecting Bax activity in mitochondria during apoptosis. They show with sub-nm spatial resolution how Bax attacks its target mitochondrial membrane, ranging from basic association to final membrane perforation by generation of pores with the removed lipids deposited in large Bax/lipid clusters above the membrane. Our comprehensive model clearly reveals how apoptotic Bcl-2 proteins such as Bax or Bak remodel the supramolecular architecture of the mitochondrial outer membrane shell to drive apoptosis further towards its ultimate goal, the removal of doomed cells.

## Materials and Methods

### Material

#### Preparation of protonated (h-Bax) and deuterated (d-Bax) protein

The methods for expression and purification of Bax was accomplished by using previous procedures (40). Deuteration (> 90%) of Bax was performed by using the following media recipe: 1 liter of M9 media containing 13 g KH_2_PO_4_, 10 g K_2_HPO_4_, 9 g Na_2_HPO_4_, 2.4 g K_2_SO_4_, 2 g NH_4_Cl, 2.5 ml of MgCl_2_ (2.5 M stock), 1 ml thiamine (30 mg/ml stock), 2 g glucose (non-deuterated), and 2 g NH_4_Cl (non-deuterated), supplemented with trace elements (1 ml of 50 mM FeCl_3_, 20 mM CaCl_2_, 10 mM MnCl_2_, 10 mM ZnSO_4_, 2 mM CoCl_2_, 2 mM CuCl_2_, 2 mM NiCl_2_, 2 mM Na_2_MoO_4_ and 2 mM H_3_BO_3_ per liter medium), 100 μg/ml carbencillin and 34 μg/ml chloramphenicol. The media was adjusted to pH 6.9 and sterile-filtered before use.

## Methods

**Neutron Reflectometry (NR) Measurements** were performed on the white beam SURF (41) and OFFSPEC (42) reflectometers at the ISIS Neutron and Muon Source (Rutherford Appleton Laboratory, Oxfordshire, UK), which use neutron wavelengths from 0.5 to 7 Å and 1 to 12 Å respectively. The reflected intensity was measured at glancing angles of 0.35°, 0.65° and 1.5° for SURF and 0.7° and 2.0° for OFFSPEC. Reflectivity was measured as a function of the wave vector transfer, Q_z_ (Q_z_ = (4π sin θ)/λ where λ is wavelength and θ is the incident angle). Data was obtained at a nominal resolution (dQ/Q) of 4.0%. The total illuminated sample length was ∼60 mm on all instruments. Measurement times for a single reflectometry data set (∼0.01 to 0.3 Å^-2^) was 40 to 60 minutes. Data collection times for kinetic data sets varied and can be seen as the x-error bar on Fig 3*D* (inset) and Fig 4*D*.

Details of the solid-liquid flow cell and liquid chromatography setup used in the experiments described here are described by us previously (43). Briefly, solid liquid flow cells containing piranha acid cleaned silicon substrates (15 mm × 50 mm × 80 mm with one 50 mm × 80 mm polished to 3 Å root mean squared roughness) were placed onto the instrument sample position and connected to instrument controlled HPLC pumps (Knauer Smartline 1000) which enabled programmable control of the change of solution isotopic contrast in the flow cell. The samples were aligned parallel to the incoming neutron beam with the beam width and height defined using two collimating slits prior to the sample position. The sample height was aligned in such a way that the neutron beam was centred on the middle of the sample surface.

### Lipid Membrane Deposition

Initially, the clean silicon substrates were characterized by NR in D_2_O and H_2_O buffer solutions. Then freshly sonicated lipid vesicle solutions (0.2 mg/ml) were introduced into the cells in the experiment buffer solution of 20 mM Tris pD/H 7.2 150 mM NaCl 2 mM CaCl_2_ and the samples incubated at 37±1°C for ∼30 min before the non-surface bound vesicles were removed by flushing the cells with 15 ml (∼5 cell volumes) of the same buffer solution before a solution of pure D_2_O was flushed into the cell. The change in osmotic gradient across the lipid vesicles bound at the solid-liquid interface caused these to rupture, thereby forming high quality (i.e. high coverage) supported lipid bilayers (SLBs) at the solid/liquid interface. The resulting bilayers were characterized by NR under three solution isotopic contrast conditions being D_2_O, Silicon Matched Water (Si-MW, 38% D_2_O) and H_2_O buffer solutions.

### Bax Interaction

Once characterisation of the SLB was complete the samples surface was placed in the correct solution isotopic contrast (D_2_O for h-proteins and H_2_O for d-proteins) and ∼6ml of a 0.1mg/ml Bax solution was injected into the flow cell (the cell volume is 3 ml) either by hand (SURF) or using a syringe pump (OFFSPEC, AL1000-220, World Precision Instruments, Hitchin, UK). In most cases the interaction of the protein with the SLB was monitored by NR with data sets collected continuously until an equilibrium interaction between the protein and the SLB was verified by no further changes in the data being observed against time. At this point a final equilibrium data set was collected and then the excess protein was flushed from the cell and the structure of the surface protein-lipid complex was examined by NR under three solution isotopic contrast conditions (D_2_O, Si-MW and H_2_O). It should be noted no difference was found in any sample between the equilibrium Bax bound data before and after flushing of the excess protein suggesting the protein-lipid complexes formed at the sample surface were stable.

### NR Data Analysis

NR data was analysed using the RasCal software (A. Hughes, ISIS Spallation Neutron Source, Rutherford Appleton Laboratory) which employs optical matrix formalism (44) to fit layered models of the structure across bulk interfaces and allows for the simultaneous analysis of multiple NR data sets collected under different sample and isotopic contrast conditions to be fully or partially constrained to the same surface structure in terms of thickness profile but vary in terms neutron scattering length density (SLD).

The NR experiments were conducted in four stages, 1, the analysis of the bare silicon surfaces, 2, the analysis of the SLB before Bax interaction, 3, analysis of the kinetics while Bax was interacting with the membrane using a single solution isotopic contrast and, finally, 4, the analysis of the equilibrium protein-lipid surface complex. For details of NR analysis see SI appendix, Methods.

## Acknowledgments

GG acknowledges financial support from the Swedish Research Council, the Kempe Foundation, the Knut and Alice Wallenberg foundation, the SciLifeLab, SwedNMR and Umeå Insamlingsstiftelse. This work was supported by ISIS beam time awards 2210172, 2010295 and 1919323.

## Data Accessibility

NR data and custom model scripts used in NR data fitting are available via https://doi.org/10.5281/zenodo.7214794. ATR-FTIR data is available by request.

## References

1. J. C. Martinou, R. J. Youle, Mitochondria in apoptosis: Bcl-2 family members and mitochondrial dynamics. Dev. Cell 21, 92–101 (2011).

2. J. Kale, E. J. Osterlund, D. W. Andrews, BCL-2 family proteins: changing partners in the dance towards death. Cell Death Differ. 25, 65–80 (2018).

3. K. J. Campbell, S. W. G. Tait, Targeting BCL-2 regulated apoptosis in cancer. Open Biol. 8, 180002 (2018).

4. S. Grabow et al., Subtle Changes in the Levels of BCL-2 Proteins Cause Severe Craniofacial Abnormalities. Cell Rep 24, 3285–3295 (2018).

5. R. B. Hill, K. R. MacKenzie, M. C. Harwig, The Tail-End Is Only the Beginning: NMR Study Reveals a Membrane-Bound State of BCL-XL. J. Mol. Biol. 427, 2257–2261 (2015).

6. G. C. Brito, W. Schormann, S. K. Gidda, R. T. Mullen, D. W. Andrews, Genome-wide analysis of Homo sapiens, Arabidopsis thaliana, and Saccharomyces cerevisiae reveals novel attributes of tail-anchored membrane proteins. BMC genomics 20, 835 (2019).

7. A. U. Mushtaq et al., Neutron reflectometry and NMR spectroscopy of full-length Bcl-2 protein reveal its membrane localization and conformation. Commun. Biol. 4, 507 (2021).

8. M. Suzuki, R. J. Youle, N. Tjandra, Structure of Bax: coregulation of dimer formation and intracellular localization. Cell 103, 645–654 (2000).

9. S. Bleicken, G. Hofhaus, B. Ugarte-Uribe, R. Schroder, A. J. Garcia-Saez, cBid, Bax and Bcl-xL exhibit opposite membrane remodeling activities. Cell Death Dis. 7, e2121 (2016).

10. A. Shamas-Din et al., Distinct lipid effects on tBid and Bim activation of membrane permeabilization by pro-apoptotic Bax. Biochem. J. 467, 495–505 (2015).

11. D. Westphal et al., Apoptotic pore formation is associated with in-plane insertion of Bak or Bax central helices into the mitochondrial outer membrane. Proc. Natl. Acad. Sci. USA 111, E4076–E4085 (2014).

12. F. Gonzalvez, E. Gottlieb, Cardiolipin: setting the beat of apoptosis. Apoptosis 12, 877–885 (2007).

13. Y. C. Lai et al., The role of cardiolipin in promoting the membrane pore-forming activity of BAX oligomers. Biochim. Biophys. Acta Biomembr. 1861, 268–280 (2019).

14. R. Salvador-Gallego et al., Bax assembly into rings and arcs in apoptotic mitochondria is linked to membrane pores. EMBO J. 35, 389–401 (2016).

15. M. Z. Zhang, J. Zheng, R. Nussinov, B. Y. Ma, Release of Cytochrome C from Bax Pores at the Mitochondrial Membrane. Sci. Rep. 7, 2635 (2017).

16. S. Bleicken et al., Topology of active, membrane-embedded Bax in the context of a toroidal pore. Cell Death Differ. 25, 1717–1731 (2018).

17. K. Cosentino et al., The interplay between BAX and BAK tunes apoptotic pore growth to control mitochondrial-DNA-mediated inflammation. Mol. Cell 82, 933–949 (2022).

18. V. Vasquez-Montes, M. V. Rodnin, A. Kyrychenko, A. S. Ladokhin, Lipids modulate the BH3-independent membrane targeting and activation of BAX and Bcl-xL. Proc. Natl. Acad. Sci. USA 118, e2025834118(2021).

19. N. Paracini, L. A. Clifton, M. W. A. Skoda, J. H. Lakey, Liquid crystalline bacterial outer membranes are critical for antibiotic susceptibility. Proc. Natl. Acad. Sci. USA 115, E7587–E7594 (2018).

20. L. A. Clifton et al., An Accurate In Vitro Model of the E. coli Envelope. Angew Chem Int Edit 54, 11952–11955 (2015).

21. H. P. Wacklin et al., Neutron reflection study of the interaction of the eukaryotic pore-forming actinoporin equinatoxin II with lipid membranes reveals intermediate states in pore formation. Biochim. Biophys. Acta Biomembr. 1858, 640–652 (2016).

22. H. N. Gong et al., Structural Disruptions of the Outer Membranes of Gram-Negative Bacteria by Rationally Designed Amphiphilic Antimicrobial Peptides. ACS Appl. Mater 13, 16062–16074 (2021).

23. L. A. Clifton et al., Design and use of model membranes to study biomolecular interactions using complementary surface-sensitive techniques. Adv Colloid Interface 277, 1012118 (2020).

24. J. D. Gelles, J. E. Chipuk, Robust high-throughput kinetic analysis of apoptosis with real-time high-content live-cell imaging. Cell Death Dis. 7, e2493 (2016).

25. D. Ardail et al., Mitochondrial contact sites. Lipid composition and dynamics. J. Biol. Chem. 265, 18797–18802 (1990).

26. S. Qian, W. C. Wang, L. Yang, H. W. Huang, Structure of transmembrane pore induced by Bax-derived peptide: Evidence for lipidic pores. Proc. Natl. Acad. Sci. USA 105, 17379–17383 (2008).

27. A. Y. Robin et al., Ensemble Properties of Bax Determine Its Function. Structure 26, 1346-+ (2018).

28. F. J. Lv et al., An amphipathic Bax core dimer forms part of the apoptotic pore wall in the mitochondrial membrane. EMBO J. 40, e106438 (2021).

29. N. R. Ader et al., Molecular and topological reorganizations in mitochondrial architecture interplay during Bax-mediated steps of apoptosis. Elife 8, e40712 (2019).

30. R. T. Uren, S. Iyer, R. M. Kluck, Pore formation by dimeric Bak and Bax: an unusual pore? Philos T R Soc B 372 (2017).

31. M. T. F. Telling et al., The dynamic landscape in the multi-subunit protein, apoferritin, as probed by high energy resolution neutron spectroscopy. Eur. Biophys. J. Biophys. Lett. 40, 156–156 (2011).

32. S. A. Tatulian, FTIR Analysis of Proteins and Protein-Membrane Interactions. Methods Mol. Biol. 2003, 281–325 (2019).

33. A. D. Cowan et al., BAK core dimers bind lipids and can be bridged by them. Nat. Struct. Mol. Biol. 27, 1024-+ (2020).

34. L. Galluzzi et al., Molecular mechanisms of cell death: recommendations of the Nomenclature Committee on Cell Death 2018. Cell Death Differ. 25, 486–541 (2018).

35. M. Lidman et al., The oxidized phospholipid PazePC promotes permeabilization of mitochondrial membranes by Bax. Biochim. Biophys. Acta Biomembr. 1858, 1288–1297 (2016).

36. A. Poulaki, S. Giannouli, Mitochondrial Lipids: From Membrane Organization to Apoptotic Facilitation. Int. J. Mol. Sci 23, 3738 (2022).

37. L. A. Clifton, Unravelling the structural complexity of protein-lipid interactions with neutron reflectometry. Biochem Soc T 49, 1537–1546 (2021).

38. D. F. Suen, K. L. Norris, R. J. Youle, Mitochondrial dynamics and apoptosis. Genes Dev. 22, 1577–1590 (2008).

39. M. A. Sani, O. Keech, P. Gardestrom, E. J. Dufourc, G. Grobner, Magic-angle phosphorus NMR of functional mitochondria: in situ monitoring of lipid response under apoptotic-like stress. FASEB J. 23, 2872–2878 (2009).

40. A. P. G. Dingeldein et al., Bax to the future - A novel, high-yielding approach for purification and expression of full-length Bax protein for structural studies. Protein Expr. Purif. 158, 20–26 (2019).

41. J. Penfold et al., Recent advances in the study of chemical surfaces and interfaces by specular neutron reflection. J. Chem. Soc. Faraday T 93, 3899–3917 (1997).

42. J. R. P. Webster, S. Langridge, R. M. Dalgliesh, T. R. Charlton, Reflectometry techniques on the Second Target Station at ISIS: Methods and science. Eur. Phys. J. Plus 126 (2011).

43. L. A. Clifton et al., The Effect of Lipopolysaccharide Core Oligosaccharide Size on the Electrostatic Binding of Antimicrobial Proteins to Models of the Gram Negative Bacterial Outer Membrane. Langmuir 32, 3485–3494 (2016).

44. M. Born, E. Wolf, A. B. Bhatia, Principles of optics: electromagnetic theory of propagation, interference, and diffraction of light (Cambridge University Press, Cambridge, 7th edition, 952 pp. 2019.

