## Supplementary Information for "Bax Forms a Membrane Surface Protein-Lipid Complex as it Initiates Apoptosis"

Gerhard Gröbner

**This PDF file includes:**

- Supporting text
- Figures S1 to S7
- Tables S1 to S3
- SI References

### Supporting Information Text

### Extended Methods

#### Analysis of NR experiments

Data was fitted using RasCal's custom model option where a bespoke script is used to describe the relationship between the fitted experimental parameters, the interfacial layers, the layer structure and the resulting SLD profile which is used to calculate the model reflectivity data sets which were fitted the experimental data. Models were constructed which constrained steps 1-4 of the experiment process to the same substrate (silicon and silicon-dioxide) structure.

For the majority of samples the membrane structure prior to Bax interaction was fitted as a four layer model, which, moving from the substrate to the bulk solution, was a thin $SiO_2$ layer, the inner bilayer head-groups, bilayer tails and the outer bilayer head-groups. The expected molecular volume and scattering length of the bilayer head/tails was determined from the molecular components of each lipid and the ratio of POPC and cardiolipin in the bilayers. The molar ratio of head and tail components in the bilayer was maintained by fitting an average lipid area per molecule for the bilayer where the thickness of the layers is defined by:

$$Layer\ thickness\ [\text{Å}] = \frac{Component\ Molecular\ Volume\ [\text{Å}^3]}{Area\ per\ molecule\ [\text{Å}^2]} \qquad \text{Eq.1}$$

And the SLD is defined by:

$$SLD\ [\text{Å}^{-2}] = \frac{\sum b\ [\text{Å}]}{Component\ Molecular\ Volume\ [\text{Å}^3]} \qquad \text{Eq. 2}$$

To allow for the difference in hydration between the hydrophobic head-groups and the hydrophilic tails of the bilayer two water parameters were fitted, membrane defect water which is hydration found both in the head-groups and tails due to defect in the SLB in the plane of the surface and head-group bound waters. In the case of all SLBs described in this study there was very little defect water detected prior to the interaction of Bax with the coverage of the SLBs on the substrate surface always approaching 100%.

Upon the equilibrium binding of Bax a number of increasingly complex models were used to satisfactorily fit the experimental data to a layer model. Ultimately, the model which suitably resolved all the features present in the experimental data sets was a seven layer model of a disrupted SLB and a protein-lipid complex on the bilayer surface. The initial four layers of the structure were the same as for the bilayer prior to the proteins interaction but with a large increase in the protein and water content and decrease in the lipid content. The three additional layers were found to be composed of a mixture of protein and lipid with a protein only layer bound to the SLB outer head-group region, a complex protein-lipid layer next to this and a protein only layer bound to this adjacent to the bulk solution.

Data sets which were collected against time during the interaction of Bax with the SLBs were fitted as a weighted average of the structures before and after the equilibrium interaction of Bax with the model mitochondrial surface.

***NR Error Estimation and Plotting:*** Bayesian inference of the ambiguity of the resolved structures from NR model-to-data fitting was undertaken using MCMCStat (https://mjlaine.github.io/mcmcstat) Delayed-Rejection Adaptive Metropolis algorithms (DRAM) (1) Monte-Carlo-Markov Chain (MCMC) routines to refit the data using a user defined number of steps. To fit data using this approach, the likelihood function is defined in terms of the Chi-squared goodness of fit criteria, as shown previously (2). The parameter uncertainties were then determined from the posterior distributions as the shortest percentile confidence interval (3) from each (in this case 65%) and the uncertainties on the reflectivity's and SLD's were generated by randomly sampling (in this case 1000 samples) from the Markov chains, calculating reflectivity's and SLD's for each set of samples, and taking the relevant percentile across all the resampled reflectivity or SLD curves at each point in $Q_z$ (or distance) to represent the uncertainties on the fits. The chain samples are also used to generate the line shading used to denote the ambiguity of the structure across the interface in the volume fraction profiles. Best fit lines are the mean of the reflectivity and SLD uncertainties and are shown as a darker line in figures.

Volume fraction profiles detailing the distribution of components across the solid/liquid interface before and after the equilibrium interaction of Bax were produced using a bespoke script. MCMC Bayesian error estimation results and the relationship between the fitting parameters and interfacial structure in the RasCal custom model were used to determine the distribution of each structural component in the volume fraction vs. distance profile. The volume fraction of an individual component was calculated in 1 Å increments across the solid/liquid interface (the silicon/silicon dioxide interface set as zero). The mean, lower, and upper 65% confidence interval bounds of each component distribution were determined for every 1 Å segment using the MCMC error estimation results or derived parameters; these confidence intervals were then used to produce a line width error region above and below the mean values. The water distribution was calculated as the remaining unoccupied volume for each 1 Å slice and summed across the interface with the appropriate error propagation.

***Attenuated Total Reflection Fourier Transform Infra-Red Spectroscopy (ATR-FTIR):*** Trapezoidal silicon substrates for attenuated total reflection infrared spectroscopy were obtained from Crystran (Poole, UK). These substrates were made to fit into a Specac (Orpington, UK) liquids ATR accessory which was, itself, fitted into the sample cavity Thermo-Fisher iS50 Infra-red spectrometer (Waltham, MA, USA). The substrates have four polished faces (to ~6 Å root mean squared roughness) the largest being a 72 mm × 10 mm face which was used as the sample surface. The IR beam enters the substrate through a polished face at 45° relative to the sample surface, total internal reflectance of the IR beam inside the substrate gives rise to six evanescent waves on the sample surface.

To reduce the influence of water vapour on the IR spectra the iS50 instrument and ATR mirror assembly accessory was continuously purged with dry air from a Peak scientific (Glasgow, UK) $CO_2$ and water removing air purge. The Specac liquid ATR cell was modified to fit Omni-fit tubing which was connected to a syringe pump (AL1000-220, World Precision Instruments, Hitchin, UK). The experimental $D_2O$ buffer solution (20 mM Tris pD 7.2 150 mM NaCl 2 mM $CaCl_2$) was injected into the ATR flow cell which was then heated to 37±1°C using a water bath (Julabo, Seelbach Germany). $D_2O$ solutions were used for all ATR-FTIR measurements as the $D_2O$ bending mode is lower (1215 cm-1) compared to $H_2O$ (1645 cm-1) meaning limited contamination of protein amide I region by spectral bands from water (4). After buffer flushing a background spectra was collected. Spectra were then collected monitoring the removal of water vapour from the spectrometer and mirror assembly until a steady state was reached. A 2nd background measurement was then collected before deposition of the SLB.

SLB deposition was conducted using the same methodology as described for NR measurements (osmotic shock) and monitored through the appearance of $CH_3$ asymmetric, $CH_2$ asymmetric and $CH_2$ symmetric stretches from the lipid tails at ~2950 $cm^{-1}$, 2920 $cm^{-1}$ and 2850 $cm^{-1}$ respectively and the appearance of a lipid carbonyl stretch at 1730 $cm^{-1}$. Once the SLB was formed BAX was injected into the solid/liquid flow cell its accumulation at the near surface region was monitored through the appearance of a protein Amide I peak.

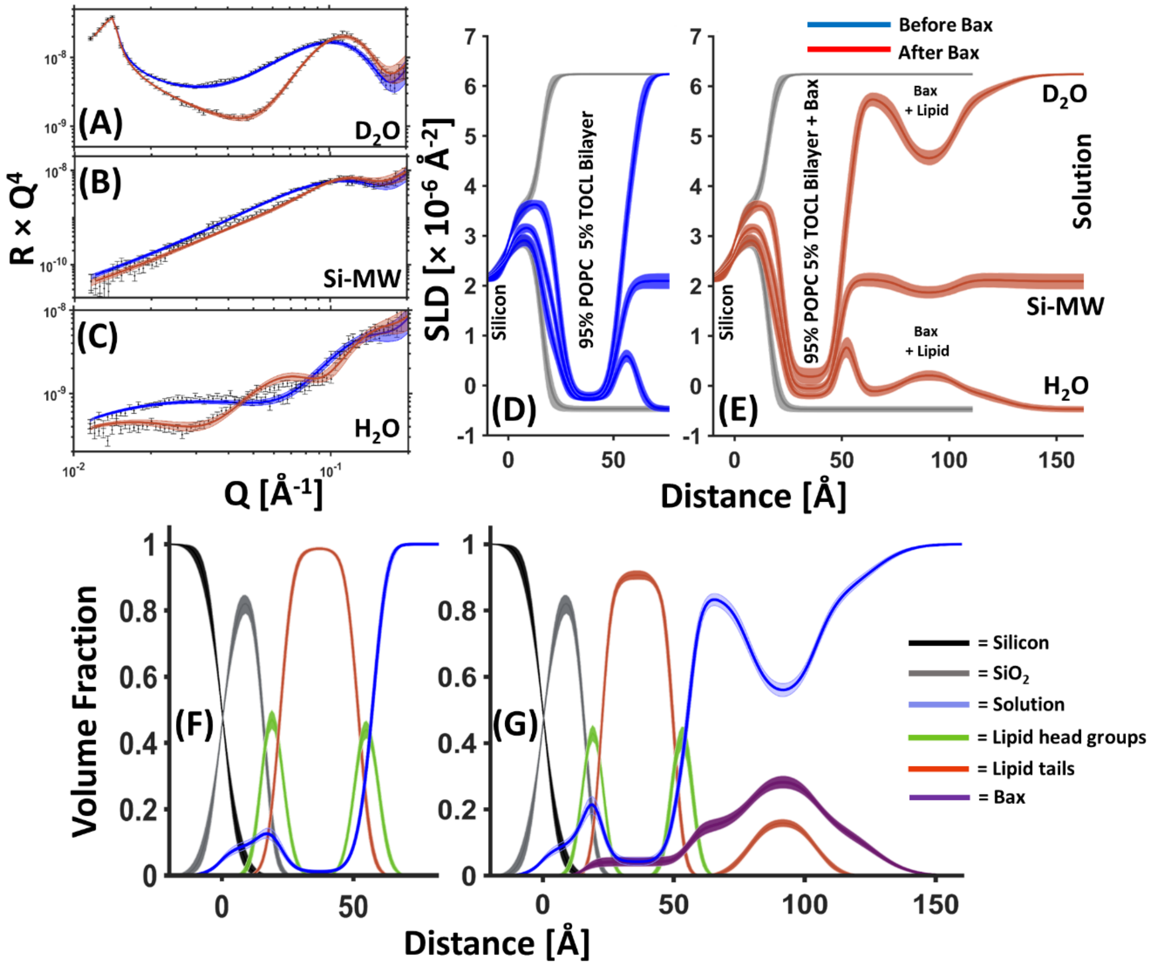


**Figure S1. NR data showing the structure at a silicon/water interface containing a POPC-5 mol% TOCL SLB before and after the interaction of Bax.** NR data (error bars) and model data fits (lines) from a POPC:TOCL SLB before (blue) and after (red) the interaction of natural abundance hydrogen (h-)Bax are shown in three differing solution isotopic contrast conditions being $D_2O$ (A), Si-MW (B) and $H_2O$ (C) buffer solutions. The scattering length density (SLD) profiles the model fits represent are shown for the surface structure before (D) and after the h-Bax interaction (E). The component volume fraction profiles at the Si/water interface before (F) and after (G) the h-Bax interaction determined from the NR fits are shown. Here the silicon distribution is given in black, silicon dioxide is grey, water is blue, lipid head-groups are green, lipid tails are red and the Bax protein distribution is in purple. Line widths in the NR data fits represent the 65% confidence interval of the range of acceptable fits determined from MCMC error analysis and the line widths on the SLD and volume fraction profiles represent the ambiguity in the resolved interfacial structure determined from these.

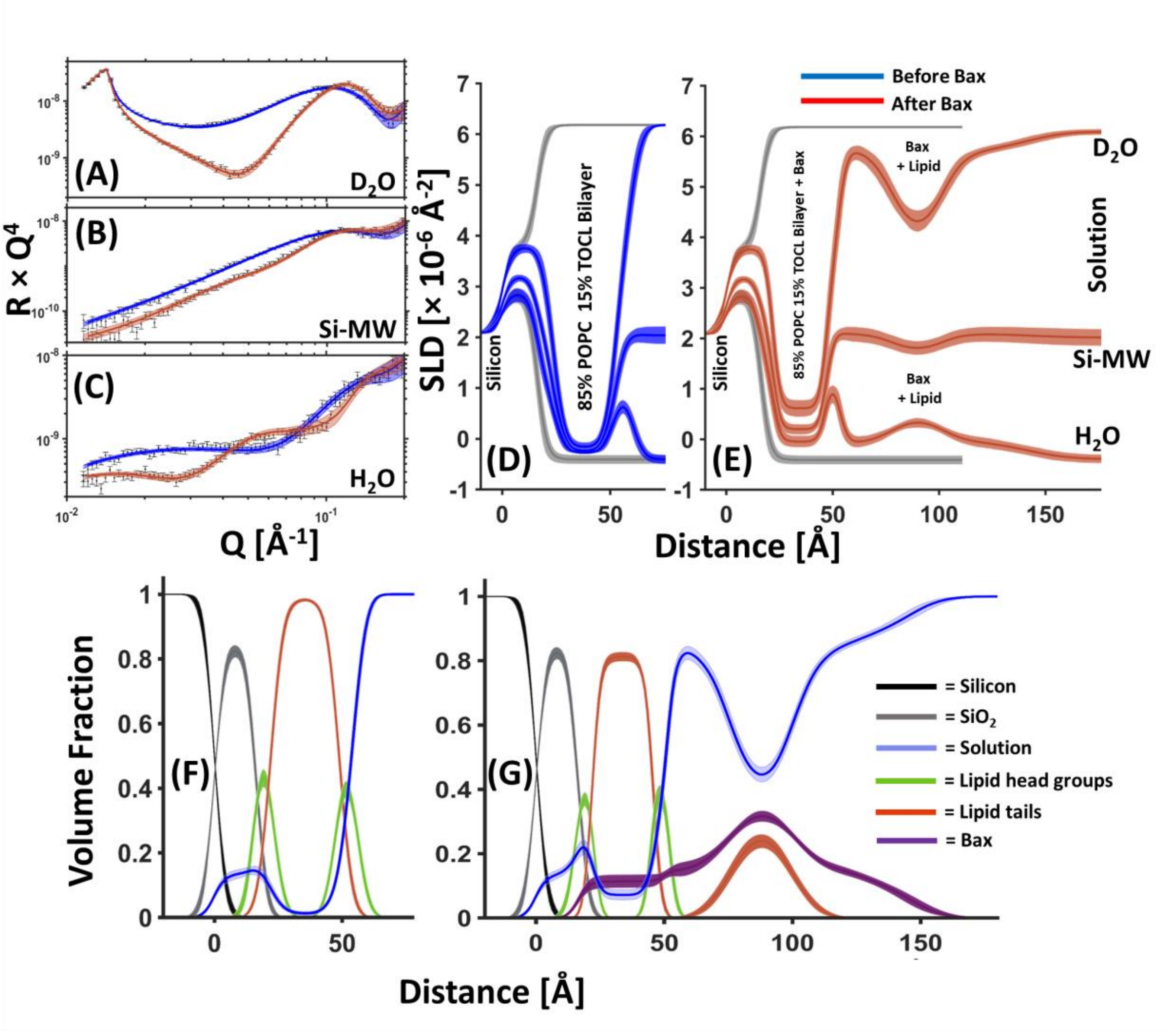


**Figure S2. NR data showing the structure at a silicon/water interface containing a POPC-15 mol% TOCL SLB before and after the interaction of Bax.** NR data (error bars) and model data fits (lines) from a POPC:TOCL SLB before (blue) and after (red) the interaction of natural abundance hydrogen (h-)Bax are shown in three differing solution isotopic contrast conditions being $D_2O$ (A), Si-MW (B) and $H_2O$ (C) buffer solutions. The scattering length density (SLD) profiles the model fits represent are shown for the surface structure before (D) and after the h-Bax interaction (E). The component volume fraction profiles at the Si/water interface before (F) and after (G) the h-Bax interaction determined from the NR fits are shown. Here the silicon distribution is given in black, silicon dioxide is grey, water is blue, lipid head-groups are green, lipid tails are red and the Bax protein distribution is in purple. Line widths in the NR data fits represent the 65% confidence interval of the range of acceptable fits determined from MCMC error analysis and the line widths on the SLD and volume fraction profiles represent the ambiguity in the resolved interfacial structure determined from these.

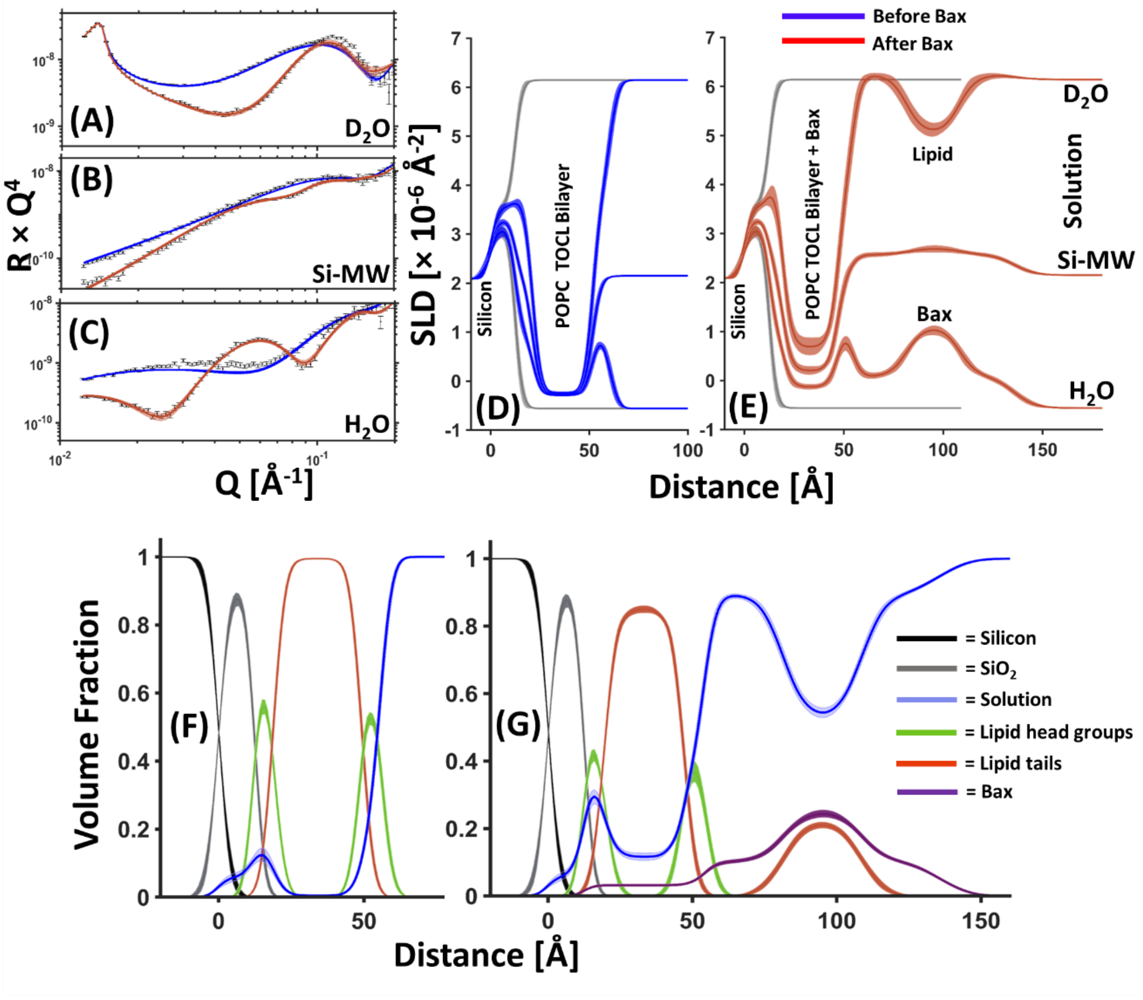


**Figure S3. NR data showing the structure at a silicon/water interface containing a POPC-10 mol% TOCL SLB before and after the interaction of deuterated (d-)Bax.** NR data (error bars) and model data fits (lines) from a POPC:TOCL SLB before (blue) and after (red) the interaction of natural abundance hydrogen (d-)Bax are shown in three differing solution isotopic contrast conditions being $D_2O$ (A), Si-MW (B) and $H_2O$ (C) buffer solutions. The scattering length density (SLD) profiles the model fits represent are shown for the surface structure before (D) and after the d-Bax interaction (E). The component volume fraction profiles at the Si/water interface before (F) and after (G) the d-Bax interaction determined from the NR fits are shown. Here the silicon distribution is given in black, silicon dioxide is grey, water is blue, lipid head-groups are green, lipid tails are red and the Bax protein distribution is in purple. Line widths in the NR data fits represent the 65% confidence interval of the range of acceptable fits determined from MCMC error analysis and the line widths on the SLD and volume fraction profiles represent the ambiguity in the resolved interfacial structure determined from these.

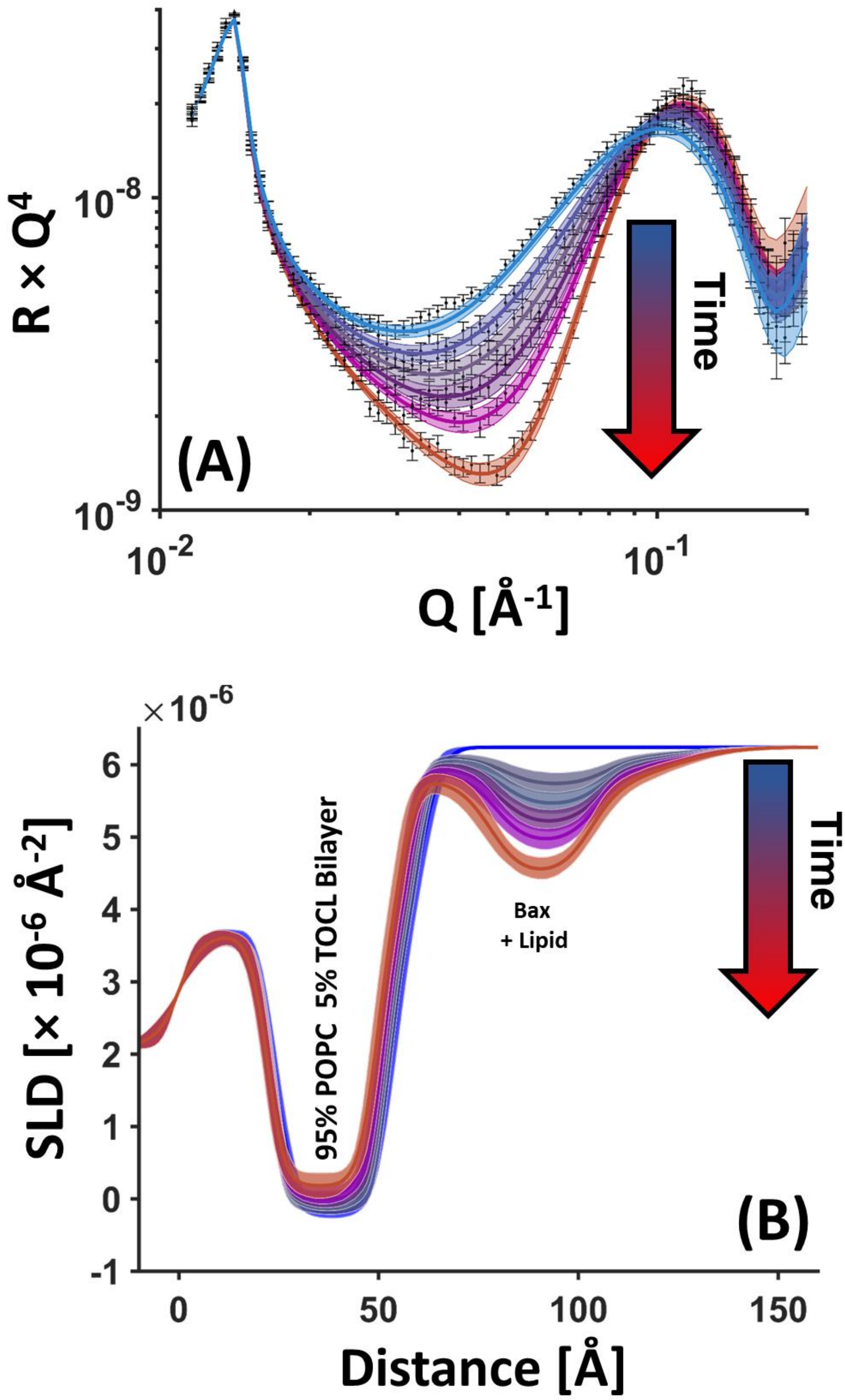


**Figure S4. Time resolved NR data (error bars) and model data fits (lines) from a POPC-5 mol% TOCL SLB.** Time dependence of NR data during the interaction of (h-)Bax in a $D_2O$ buffer solution (A); and the changes in the scattering length density of the surface assemblage against time (B, depicted from blue to red). Note how the increasing SLD of the bilayer, depicting lipid loss, is commensurate with decreasing SLD of the Bax/Lipid surface complex, which depicting increasing complex coverage. Lines widths on the fits/SLD profiles depict the ambiguity in the resolved structure from MCMC error estimation.

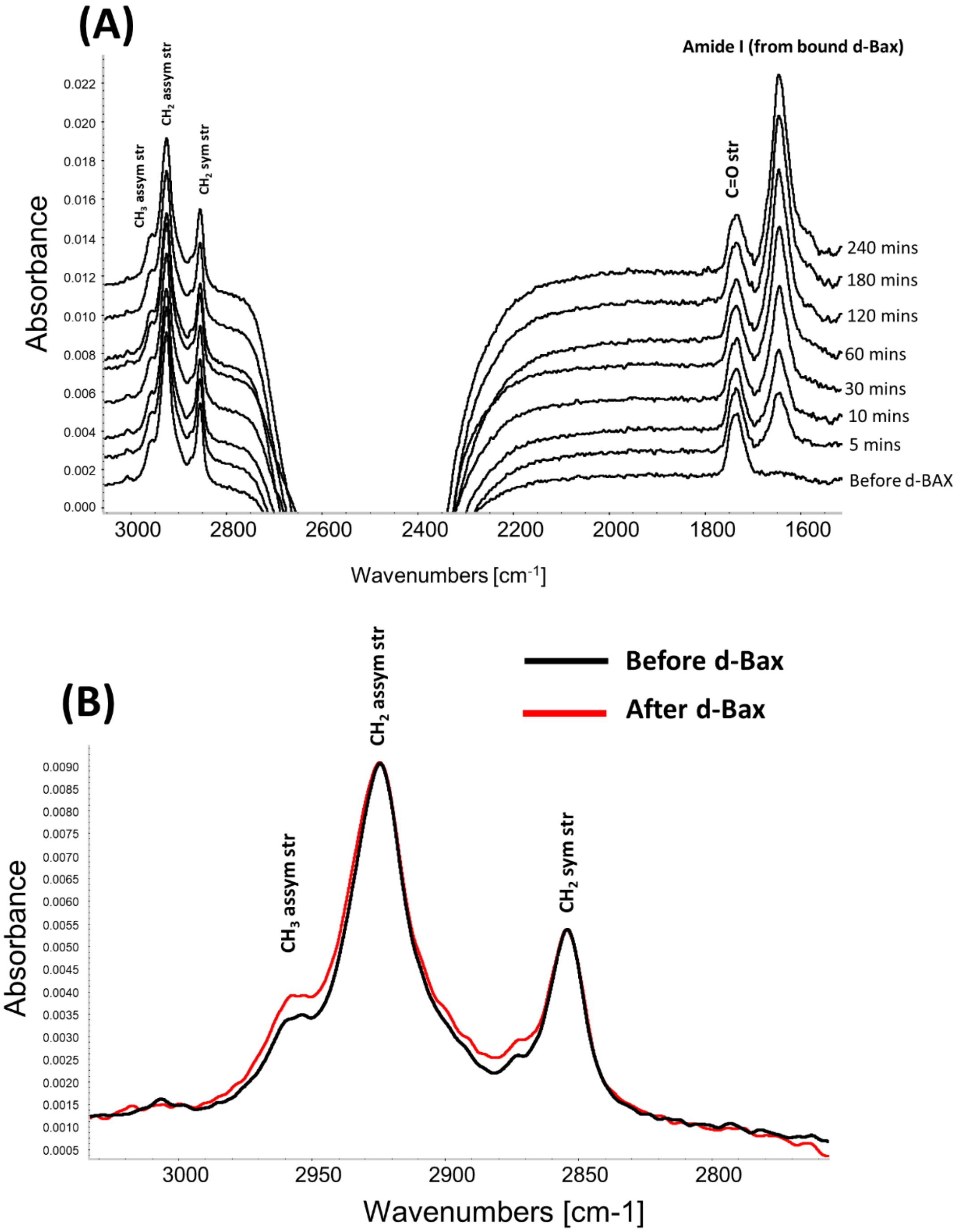


**Figure S5. ATR-FTIR spectra obtained during the interaction of d-Bax with a POPC-10 mol% TOCL SLB at the silicon/$D_2O$ buffer interface.** Time dependence of the main spectral features (A); A comparison (B) of the $CH_2$ stretch region of the IR spectra showing the differences in the bands in this region before (back line) and after equilibrium d-Bax binding (red line).

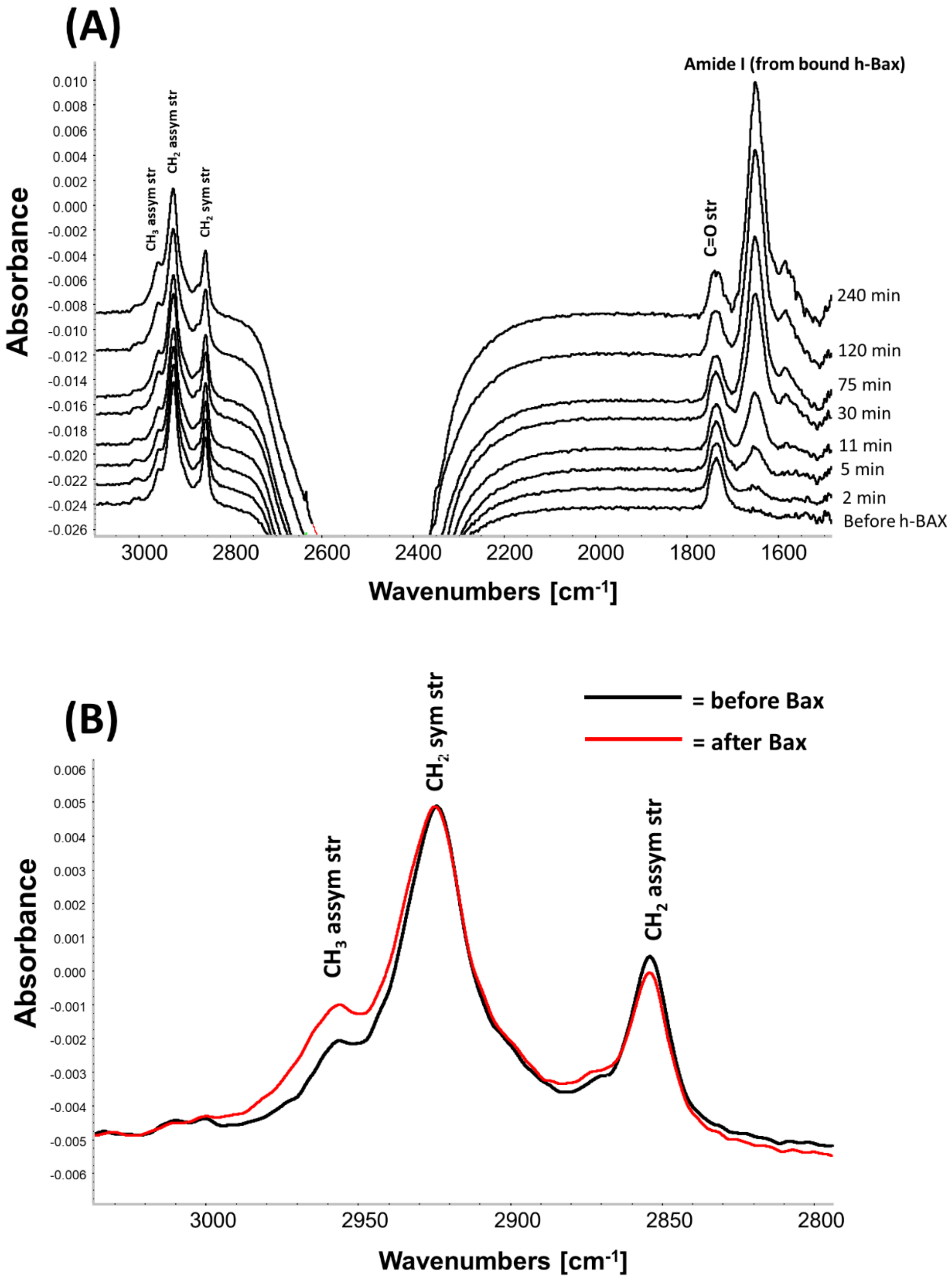


**Figure S6. ATR-FTIR spectra obtained during the interaction of h-Bax with a POPC-10 mol% TOCL SLB at the silicon/$D_2O$ buffer interface.** Time-dependence of the main spectral features (A); A comparison (B) of the $CH_2$ stretch region of the IR spectra showing the differences in the bands in this region before (back line) and after equilibrium h-Bax binding (red line).

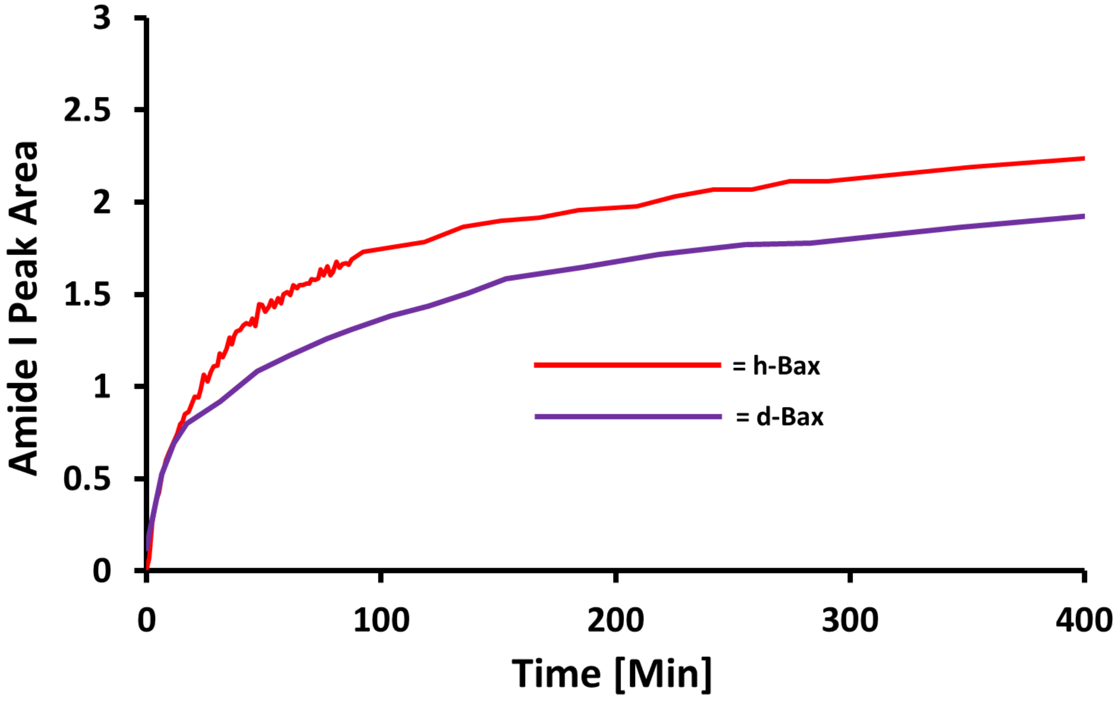


**Figure S7. A comparison between the change in Amide I peak area vs. time for h-Bax (red) and d-Bax (purple) during the interaction of these proteins with POPC-10 mol% TOCL SLBs examined by ATR-FTIR.** Examples of the spectra used to generate this plot are shown in Figure S5 for d-Bax and S6 for h-Bax. Peak areas were normalized to the $CH_2$ asymmetric stretch from the bilayer lipid tails.

**Table S1. The resolved structural components before and after the interaction of (h-)Bax with SLB composed of POPC- 5 mol% TOCL.***

| | Average Lipid Area per Molecule [Å$^2$] | Tails Thickness [Å] | Tail Composition [vol%] | Head Group Thickness [Å] | Outer head-group Composition [vol%] | Membrane Surface Bax/Lipid Complex Thickness [Å] | Bax Surface Layer Composition [vol%] |
|---|---|---|---|---|---|---|---|
| **POPC:TOCL (95:5 mol%)** | 64.8 Å$^2$ (62.4 Å$^2$, 64.2 Å$^2$) | 30.3 Å (30.0 Å , 30.6 Å) | Lipid 99% (98%, 100%)<br><br>Solution 1% (0%, 2%) | 6.2 Å (6.0 Å ,6.4 Å) | Head-group 83% (80%, 85%)<br><br>Water 17% (15%, 20%) | - | - |
| **POPC:TOCL (95:5 mol%) + h-Bax** | 70.1 Å$^2$ (69.0 Å$^2$, 71.3 Å$^2$) | 27.3 Å (26.8 Å, 27.7 Å) | Lipid 91% (90%, 92%)<br><br>Protein 4% (3%, 5%)<br><br>Solution 4% (3%, 5%) | 6.6 Å (6.2 Å, 7.0 Å) | Lipid 66% (63%, 70%)<br><br>Protein 4% (3%, 5%)<br><br>Solution 29% (26%, 32%) | **1**, 22.6 Å (21.4 Å, 23.6 Å)<br><br>**2**, 24.8 Å (23.2 Å, 26.0 Å)<br><br>**3**, , 22.6 Å (21.4 Å, 23.6 Å)<br><br>**Total : 70.4 Å (69.1 Å, 71.7 Å)** | **Protein Lipid Distribution Composition:**<br>1, Protein 15% (14%, 16%) Water 85 % (84%, 86%)<br><br>2, Protein 29 % (27%, 32%) Lipid 15% (16%, 18%) Solution 55% (52%, 57%)<br><br>3, Protein 15% (14%, 16%) Water 85 % (84%, 86%) |

* Values in brackets represent the 65% confidence intervals determined from MCMC resampling of the experimental data fits.

**Table S2. The resolved structural components before and after the interaction of (h-)Bax with SLB composed of POPC- 15 mol% TOCL .***

| | Average Lipid Area per Molecule [Å$^2$] | Tails Thickness [Å] | Tails Composition [vol%] | Head Group Thickness [Å] | Outer head-group Composition [vol%] | Membrane Surface Bax/Lipid Complex Thickness [Å] | Bax Surface Layer Composition [vol%] |
|---|---|---|---|---|---|---|---|
| **POPC:TOCL (85:15 mol%)** | 71.4 Å$^2$<br>(70.7 Å$^2$, 72.2 Å$^2$) | 26.9 Å<br>(26.6 Å, 27.2 Å) | Lipid 99%<br>(98%, 100%)<br><br>Solution 1%<br>(0%, 2%) | 5.6 Å<br>(5.4 Å ,5.8 Å) | Head-group 82%<br>(79%, 85%)<br>Water 18%<br>(15%, 21%) | - | - |
| **POPC:TOCL (85:15 mol%) + h-BAX** | 81.1 Å$^2$<br>(79.8 Å$^2$, 82.4 Å$^2$) | 23.6 Å<br>(23.3 Å, 24.0 Å) | Lipid 81%<br>(80%, 82%)<br><br>Protein 11%<br>(10%, 14%)<br><br>Solution 8%<br>(6%, 10%) | 5.7 Å<br>(5.4 Å, 6.0 Å) | Lipid 60%<br>(57%, 64%)<br><br>Protein 11%<br>(10%, 14%)<br><br>Solution 29%<br>(25%, 32%) | **1**, 24.5 Å<br>(22.6 Å, 25.9 Å)<br><br>**2**, 23.7 Å<br>(21.0 Å, 27.0 Å)<br><br>**3**, , 43.0 Å<br>(38.7 Å, 47.3 Å)<br><br>**Total : 91.7 Å**<br>**(87.3 Å, 96.3 Å)** | **Protein Lipid Distribution Composition:**<br>1, Protein 14% (11%, 17%)<br>Water 86 % (83%, 89%)<br><br>2, Protein 35 % (32%, 39%)<br>Lipid 19% (17%, 21%)<br>Solution 46% (41%, 50%)<br><br>3, Protein 15% (13%, 17%)<br>Water 85 % (83%, 87%) |

* Values in brackets represent the 65% confidence intervals determined from MCMC resampling of the experimental data fits.

**Table S3. The resolved structural components before and after the interaction of (d-)Bax with SLB composed of 9:1 (mol/mol) POPC:TOCL.***

| | Average Lipid Area per Molecule [$Å^2$] | Tails Thickness [Å] | Tails Composition [vol%] | Head Group Thickness [Å] | Outer head-group Composition [vol%] | Membrane Surface Bax/Lipid Complex Thickness [Å] | Bax Surface Layer Composition [vol%] |
|---|---|---|---|---|---|---|---|
| **POPC:TOCL (90:10 mol%)** | 63.1 $Å^2$ (62.7 $Å^2$, 63.6 $Å^2$) | 30.4 Å (30.2 Å, 30.6 Å) | Lipid 100% (99%, 100%)<br><br>Solution 0% (0%, 1%) | 6.3 Å (6.1 Å ,6.5 Å) | Head-group 84% (81%, 86%)<br><br>Water 16% (14%, 19%) | - | - |
| **POPC:TOCL (90:10 mol%) + d-BAX** | 69.4 $Å^2$ (68.4 $Å^2$, 70.4 $Å^2$) | 27.6 Å (27.2 Å, 28.0 Å) | Lipid 85% (81%, 85%)<br><br>Protein 3% (2%, 4%)<br><br>Solution 12% (11%, 13%) | 7.0 Å (6.7 Å, 7.2 Å) | Lipid 59% (57%, 61%)<br><br>Protein 3% (2%, 4%)<br><br>Solution 38% (36%, 40%) | **1**, 28.0 Å (26.9 Å, 29.0 Å)<br><br>**2**, 25.9 Å (24.2 Å, 27.9 Å)<br><br>**3**, 28.0 Å (26.9 Å, 29.0 Å)<br><br>**Total: 83.1 Å (82.0 Å, 84.2 Å)** | **Protein Lipid Distribution Composition:**<br>1, Protein 10% (9%, 11%)<br>Water 90% (89%, 91%)<br><br>2, Protein 26% (25%, 27%)<br>Lipid 22% (21%, 23%)<br>Solution 51% (49%, 53%)<br><br>3, Protein 10% (9%, 11%)<br>Water 90% (89%, 91%) |

* Values in brackets represent the 65% confidence intervals determined from MCMC resampling of the experimental data fits

### SI References


1. H. Haario, M. Laine, A. Mira, E. Saksman, DRAM: Efficient adaptive MCMC. *Stat. Comput.* **16**, 339-354 (2006).
2. D. S. Sivia, J. R. P. Webster, The Bayesian approach to reflectivity data. *Physica B* **248**, 327-337 (1998).
3. D. S. Sivia, J. Skilling, *Data analysis : a Bayesian tutorial*, Oxford science publications (Oxford University Press, Oxford ; New York, ed. 2nd, 2006), pp. xii, 246 p.
4. B. Stuart, D. J. Ando, *Biological applications of infrared spectroscopy*, Analytical Chemistry by Open Learning (Published on behalf of ACOL (University of Greenwich) by John Wiley, Chichester ; New York, 1997), pp. xx, 191 p.
